# Mitochondrial event localiser (MEL) to quantitatively describe fission, fusion and depolarisation in the three-dimensional space

**DOI:** 10.1101/2020.02.12.945451

**Authors:** Rensu P Theart, Jurgen Kriel, Ben Loos, Thomas R Niesler

## Abstract

Mitochondrial fission and fusion play an important role not only in maintaining mitochondrial homeostasis but also in preserving overall cellular viability. However, quantitative analysis based on the three-dimensional localisation of these highly dynamic mitochondrial events in the cellular context has not yet been accomplished. Moreover, it remains largely uncertain where in the mitochondrial network depolarisation is most likely to occur. We present the mitochondrial event localiser (MEL), a method that allows high-throughput, automated and deterministic localisation and quantification of mitochondrial fission, fusion and depolarisation events in large three-dimensional microscopy time-lapse sequences. In addition, MEL calculates the number of mitochondrial structures as well as their combined and average volume for each image frame in the time-lapse sequence. The mitochondrial event locations can subsequently be visualised by superposition onto the fluorescence micrograph z-stack. We apply MEL to both control samples as well as cells that have been treated with hydroxychloroquine sulphate (HCQ) and observe that fission and fusion events mostly occur around the terminal branches of the mitochondrial network, with few events detected in the strongly networked areas such as the perinuclear region. Depolarisation was most abundant in the HCQ treated samples, with the majority of the depolarisation events occurring in the cell periphery.

## Introduction

Mitochondria are highly dynamic organelles that do not operate in a stagnant or isolated manner. Rather, they function in a highly energetic and networked fashion, continuously subjected to rapid remodelling events referred to as fission (fragmentation) and fusion [1]. These events serve as a critical quality control mechanism, not only to adapt and respond to changing metabolic demands but also to enable separation of damaged mitochondria from those in an interconnected state and thereby exposing them to specific degradation [2]. The elimination of damaged mitochondria, mediated through a mitochondrial specific autophagy referred to as mitophagy, decreases the risk of mitochondrial DNA (mtDNA) mutation accumulation. It also improves the electron transport chain (ETC) efficiency [3–6]. The transport of electrons between different complexes on the mitochondrial membrane results in the maintenance of a continuous voltage across the inner mitochondrial membrane (IMM) referred to as the mitochondrial membrane potential (Δ*ψ*m) [7]. Mitochondrial dynamics are highly dependent on this membrane potential. Excessive membrane potential dissipation has, for example, been observed to render fragmented mitochondria incapable of re-fusing, thereby metabolically debilitating the cell [7–9]. Furthermore, the recruitment of the mitophagy initiation protein PTEN-induced kinase 1 (PINK1) only occurs once a certain depolarisation threshold is reached [10]. Failure to eliminate damaged mitochondria is a hallmark of neurodegenerative diseases such as Parkinson’s disease [11], while certain heritable diseases, including Charcot-Marie-Tooth Type IIA, are related to dysregulated mitochondrial dynamics [12]. This underscores the important role mitochondrial fission and fusion play, not only in maintaining mitochondrial homeostasis, but also in preserving overall cellular viability [13]. It is thus not surprising that there is a lot of interest among researchers to quantify these dynamic changes accurately.

Previous attempts to better describe and unravel the interplay between mitochondrial dynamics and cell death onset have largely focused on either the change in the mitochondrial network of the entire cell or the rate at which fission and fusion occurs [11, 14, 15]. The accurate description of the quantitative relationship between fission and fusion dynamics in a three-dimensional cellular context, in order to detect deviation from its equilibrium, has however remained challenging. Moreover, although it is becoming increasingly clear that intracellular localisation of mitochondria is indicative of regional specific functions, for example the reliance of cellular organelles on mitochondrial ATP provision, it remains largely uncertain where in the mitochondrial network depolarisation is most likely to occur. It therefore remains to be determined which areas of the mitochondrial network are preferentially depolarised to facilitate either transportation or degradation and how these areas relate to the mitochondrial morphometric parameters usually employed. Currently, the methodologies available to address this question are limited, particularly in the context of three-dimensionally-based quantification of fission and fusion dynamics concomitantly with the onset of mitochondrial depolarisation. To gain a better understanding of mitochondrial dynamics, a high-throughput, automated and deterministic method that is able to localise and quantify the number of mitochondrial events in large three-dimensional (3D) time-lapse sample sets would therefore be advantageous. Such a method would allow subsequent quantitative description of the fission/fusion equilibrium as well as the extent of depolarisation.

Using time-lapse microscopy data of cells labelled for mitochondria, it has previously been observed that fission and fusion are rapid events occurring within a five second time frame under homeostatic conditions [16]. This work, however, did not investigate mitochondrial depolarisation. Current methods typically rely on the manual comparison of two time-lapse image frames in order to observe where fission, fusion or depolarisation occurred. This very labour intensive approach makes it challenging to gain comprehensive insights into the mitochondrial dynamics of a whole three-dimensional sample. It is usually unclear whether mitochondrial fission and fusion events are changing spatio-temporally, thereby shifting the equilibrium towards either fission or fusion. In this context, it is also often not clear whether a cell with a more extensively fused mitochondrial network is in transition or whether is has established a new equilibrium between fission and fusion by means of a newly adapted net contribution of fusion events. Similarly, a cell with greater mitochondrial fragmentation may establish a new equilibrium between fission and fusion as an adaptive response, in order to maintain and preserve this fragmented morphology. Hence, the micrograph reflects an underlying relationship between fission and fusion and this relationship is currently challenging to describe. Although mitochondrial photoactivation [14, 15] provides a highly selective tool to quantitatively assess mitochondrial dynamics, it does not reveal the relative contribution of fission and fusion events to the observed dynamics. For example, it has been noted that mitochondrial fusion is often enhanced during adaptations to metabolic perturbations, particularly in the perinuclear region, and may protect mitochondria from degradation [17]. On the other hand, a degree of fragmentation is required and desirable to allow, for example, mitochondrial transport in neurons to reach synaptic connections. Yet, describing the precise occurrence of fission, fusion and depolarisation has remained challenging.

To address these current challenges, we have developed an approach that automatically localises and count the number of mitochondrial events occurring between two micrograph frames in a time-lapse sequence. The results of this automatic analysis allow the quantitative assessment of fission, fusion and depolarisation localisation with high accuracy and precision. We refer to this newly developed procedure as the *mitochondrial event localiser* (MEL), because it allows us to indicate the precise three-dimensional location at which fission, fusion and depolarisation are likely to occur next. Moreover, MEL enables the determination of the individual locations of smaller structures that fuse to form a larger central structure in the two time-lapse frames that are considered. Similarly, the locations of smaller structures that will separate from a common central structure due to fission are identified. In doing so, MEL provides a platform to better understand mitochondrial dynamics in the context of health and disease, with both screening and diagnostics potential. This approach is, as far as we are aware, the first automated method for the detection of depolarised mitochondria in the context of fission and fusion events. MEL can serve both as a standalone method or as part of the broader mitochondrial analysis pipeline to enable high-throughput analysis of time-lapse data.

## Method

The development of MEL was inspired by previous work which applies a “vote casting” methodology to individual mitochondrial structures in two consecutive time-lapse frames in order to identify the likely mitochondrial event (fission, fusion, or none) the structure will undergo [16]. The purpose of this approach was specifically to count the number of fission and fusion events and can not easily be extended to determine their location. MEL was designed from the outset to both count the number of mitochondrial events and to localise each in 3D space.

The MEL algorithm processes a fluorescence microscopy time-lapse sequence of z-stack images that were stained for mitochondria and produces the 3D locations of the mitochondrial events occurring at each time step. These locations can subsequently be superimposed on the z-stacks in order to indicate the different mitochondrial events. The algorithm has been organised into two consecutive steps, namely the image pre-processing step which normalises and prepares the time-lapse frames, and the automatic image analysis step which calculates the location of the mitochondrial events based on the normalised frames.

### Image pre-processing

The image pre-processing step receives a time-lapse sequence of z-stacks as input and begins by selecting two z-stacks, which we will refer to as Frame 1 and Frame 2, for further processing. Depending on the temporal resolution that is desired, the selected z-stacks can either be consecutive time-lapse frames, or some number *k* of intermediate frames may be skipped. Since mitochondrial movement is highly dynamic, a time interval that is too long might prevent the algorithm from accurately matching mitochondrial structures between the chosen frames, thereby degrading the ability to determine mitochondrial events. The selected Frames 1 and 2 are then each processed in the same way by the image pre-processing step to generate several new image stacks which are passed to the automatic image analysis step. The image pre-processing step, which will be described in the remainder of this section, is illustrated in Fig 1.

Since MEL is not based on the analysis of a single cell, it is not necessary to select regions of interests (ROIs) before the analysis. However, since the image acquisition parameters that are used vary widely between different time-lapse sequences, we first normalise the fluorescence intensity data of the z-stacks. This is necessary since MEL relies on an accurate binarisation of the fluorescence image stack to identify the voxels that contain mitochondria. These thresholding algorithms require images to be sharp, contain minimal noise, and have good contrast between the foreground and background. The normalisation and binarisation, which form the initial part of the MEL image pre-processing step, were guided by previous work focusing on quantitative mitochondrial network analysis [16, 18–20].

**Fig 1.**
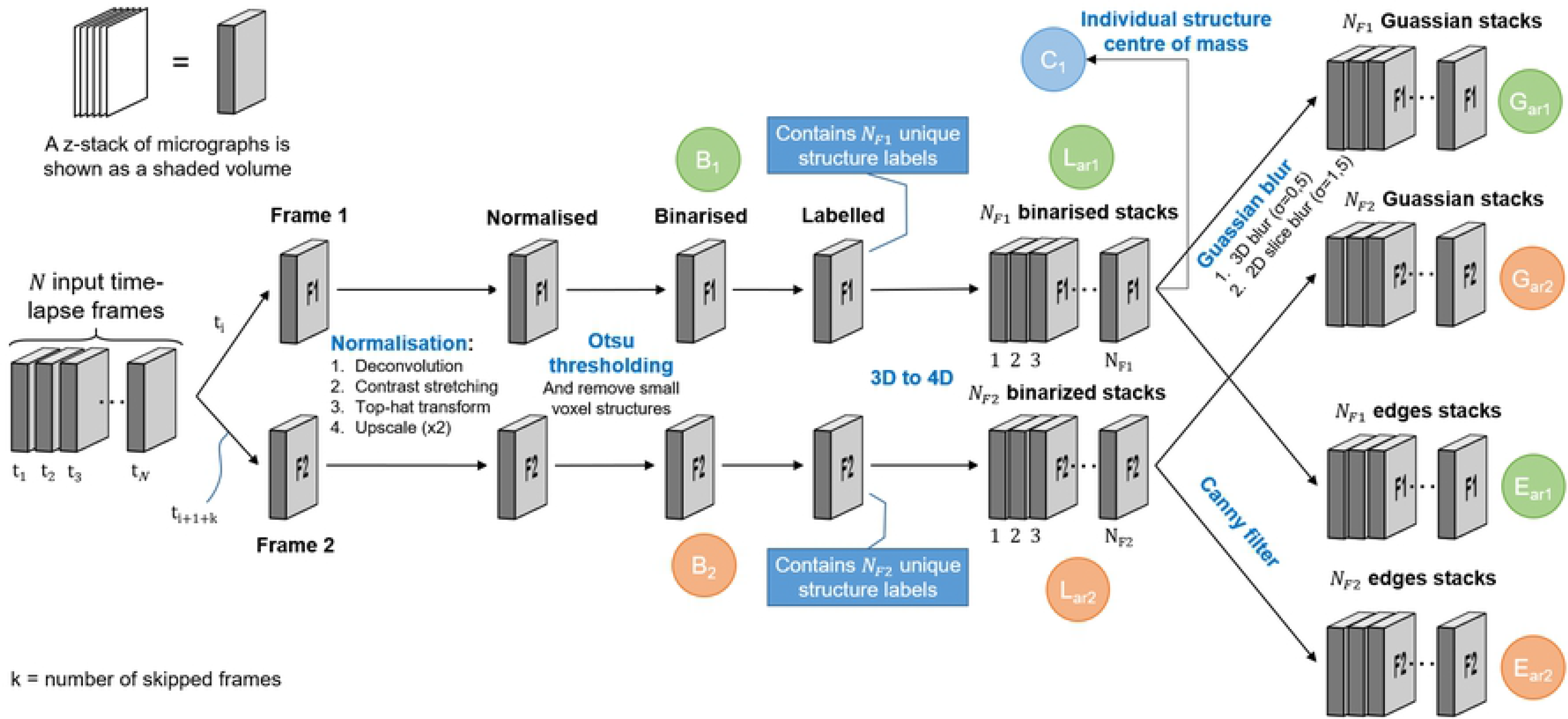
**Image pre-processing** begins by choosing two frames from the input time-lapse sequence, Frame 1 and Frame 2. Normalisation, binarisation, and labelling are then performed on both frames. The labelled frames are each separated into an array of z-stacks, where each stack contains only a single labelled structure. Each of the stacks in the array of binarised stacks, for both Frames 1 and 2, is then Gaussian blurred and Canny filtered. The latter contains only the edges of the labelled structures. The stacks that are used during the automatic image analysis step are labelled in green for Frame 1 and orange for Frame 2. Finally, an array containing the centre of mass for each labelled structure is shown in blue.

The first step in normalising the raw micrographs is to apply deconvolution to the z-stacks using a point spread function (PSF) that is estimated from the microscope’s acquisition parameters. For this, we used Huygens Professional deconvolution software. Next, we apply contrast stretching to the z-stack to normalise the fluorescence intensity between micrographs. The micrographs constituting the z-stack are then upscaled by a factor of two using bilinear interpolation so as to increase the resolution of the binarised image and reduce the possibility that two adjacent but unconnected structures are erroneously joined after binarisation. Finally, we apply a three-dimensional top-hat transformation which further reduces noise and enhances the mitochondria in such a way that they can be more easily isolated from the background.

The normalised frames are binarised by applying Otsu thresholding [21] to the z-stack, although other thresholding methods might also be suitable. Since it has been observed that some noise remnants were also being binarised, we removed any structure containing less than some appropriately small number of voxels (in our case 20) in the upscaled images. This z-stack that results from this is labelled *B* in Fig 1. Note that we differentiate between the stacks of Frame 1 and Frame 2 in Fig 1 by using subscripts 1 and 2, respectively.

After binarisation, we assign a label to each separate 3D voxel structure. The total number of labels in Frame 1 and Frame 2, as depicted in Fig 1, are *N*_*F*1_ and *N*_*F*2_, respectively. Later in the automatic image analysis step, each labelled structure in each frame will be compared with labelled structures both within the same frame as well as in the other frame. For this reason, we separate the labelled z-stack into an array of z-stacks, where the *i*^th^ stack in the array contains only the binarised voxel structure associated with label number *i*. We refer to this array of stacks as *L*_*ar*_. The subscript “*ar*” is used to indicate an *array* of stacks, containing *N*_*F*1_ and *N*_*F*2_ stacks for Frame 1 and Frame 2, respectively. This step can be thought of as adding the structure label as a fourth dimension to the data and later enables fast comparison between different labelled structures.

From the array of labelled z-stacks *L*_*ar*_ we create two similar arrays also required by the automatic image analysis step. The first is the result of applying a 3D Gaussian filter to each z-stack in the array. This slightly blurs the edges of the labelled stacks and results in the array of z-stacks *G*_*ar*_. The Gaussian blurring enhances the ability of the algorithm to match the moving mitochondrial structures between two frames by slightly inflating the structures. The second array of stacks is generated by removing all voxels that are not located on the edges of the 3D labelled structures in *L*_*ar*_ by using Canny edge detection [22]. We refer to the resulting array of z-stacks as *E*_*ar*_. This is performed mainly to improve the efficiency of the algorithm that later determines the location of fission and fusion events.

This concludes the image pre-processing step. The tuneable parameters in this step is the volume of the structures that are considered as noise, and the standard deviation with which the z-stacks are Gaussian blurred. The z-stack *B*, and the arrays of z-stacks *G*_*ar*_ and *E*_*ar*_ are now passed into the automatic image analysis step that calculates the location of the mitochondrial events.

### Automatic image analysis step

The purpose of the automatic image analysis step is to generate a list of locations in Frame 1 at which the mitochondrial events occur. It achieves this by receiving the z-stacks that were generated in the image pre-processing step. These are then automatically analysed to produce a list of the mitochondrial event locations. The automatic image analysis step is shown in Fig 2.

**Fig 2.**
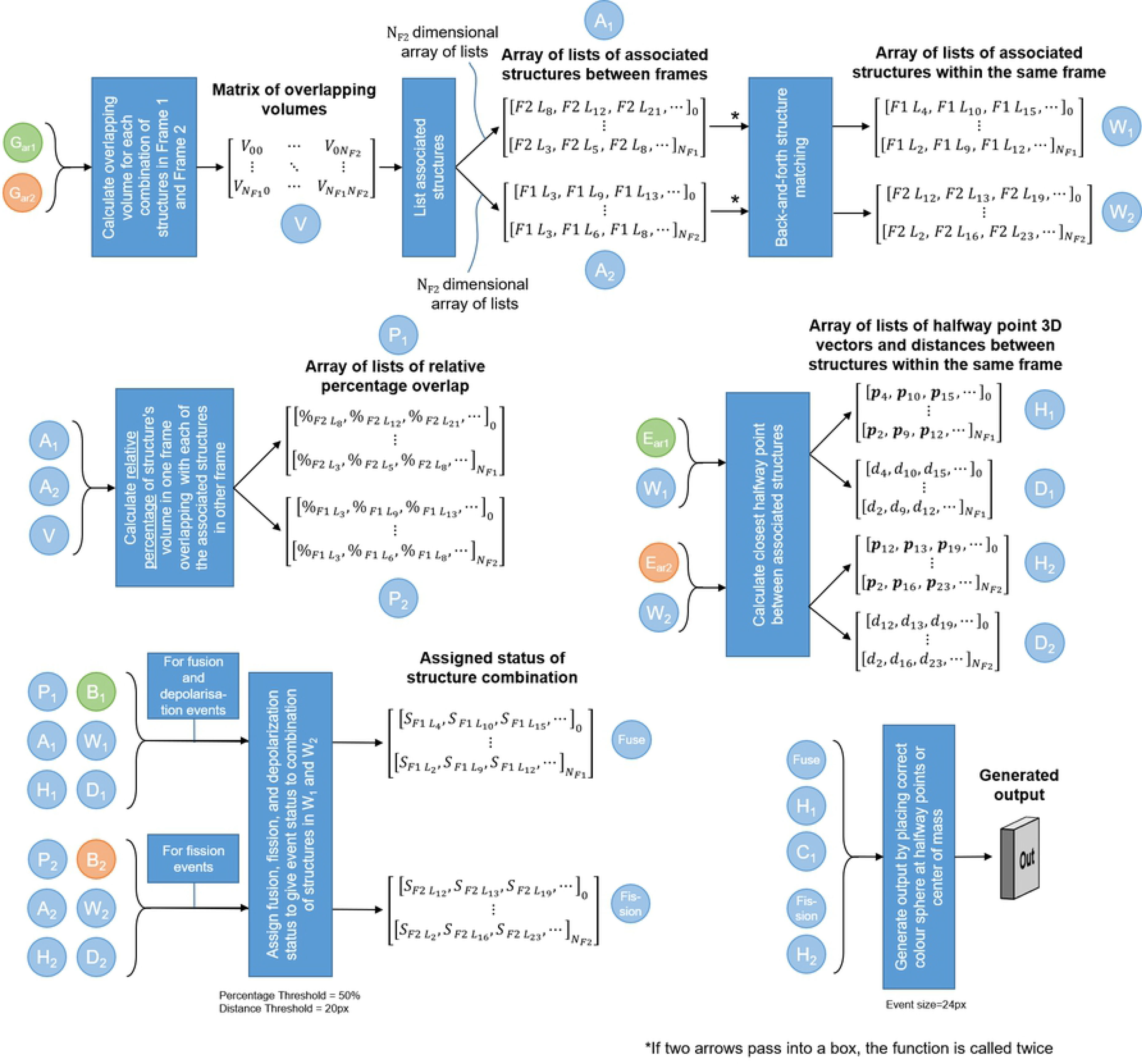
**Automatic image analysis** begins by calculating the overlapping volume of each structure in Frame 1 with each structure in Frame 2. *V*_*xy*_ refers to the number of voxels that overlap between structure *x* in Frame 1 and structure *y* in Frame 2. To compensate for such coincidental matches, all overlapping volumes that account for less than 1% of the volume of either structure in question are eliminated. Label 0 refers to the background. Using matrix *V*, the labelled structures that are associated with each other both between Frames 1 and 2, as well as within the same frame, can be calculated. The relationships are encoded as arrays of lists, where each list is related to the labelled structure corresponding to the array index. Therefore, the 8^th^ list in the array relates to labelled structure number 8. *F*2*L*_8_ refers to label number 8 in Frame 2 that is associated with the structure at the array index. Next, the relative percentage of overlap (e.g. 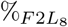) is calculated for the associated structures between the frames, and is interpreted in conjunction with *A*_1_ and *A*_2_. The midway points (e.g. **p**_4_) and distances (e.g. *d*_4_) between two structures within the same frame are calculated, and are interpreted in conjunction with *W*_1_ and *W*_2_. Using these arrays of lists, the status (e.g. 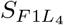) of fission, fusion, depolarisation or no event can be determined for each mitochondrial structure. Finally, the event can be overlaid on the original z-stack at the location of the midway points.

### Mitochondrial events

The three different mitochondrial events that we consider, namely fission, fusion and depolarisation, are defined in terms of changes in mitochondrial morphology that occur in the time interval between Frames 1 and 2. Specifically, fission is defined as the separation of a larger mitochondrial structure to two smaller structures. Similarly, fusion is defined as the joining of two mitochondrial structures to form a single larger structure. Finally, depolarisation is defined as the disappearance of a mitochondrial structure from Frame 1 to Frame 2 manifested by a complete loss of fluorescence signal. If more than two structures fuse to a single central structure, or if one structure undergoes fission to form more than two smaller structures, then separate locations are assigned to each.

### Process

In order to determine the location of mitochondrial events, we first identify which of the voxel structures labelled during the image pre-processing step are candidates for fission and fusion. First, the voxel structures in Frame 1 that occupy the same space as a voxel structures in Frame 2, and vice versa, are identified. For the sake of conciseness, we will refer to such colocation of structures between the two frames as overlap. Using this overlap information, we can determine which voxel structures are associated with each other within the same frame. In Frame 1, these associated structures will be the fusion candidates, and in Frame 2 they will be the fission candidates. Structures in Frame 1 that have no associated structures in Frame 2 undergo depolarisation.

To determine the associated structures between Frame 1 and Frame 2, we first calculate a matrix *V* containing the overlapping volume (in voxels) of each structure in Frame 1 with each structure in Frame 2 using the blurred arrays of z-stacks 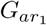 and 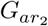. Label number 0 is reserved for the background. Blurred stacks are used in order to allow for movement of the mitochondria between frames. Using matrix *V*, we calculate two arrays *A*_1_ and *A*_2_. Each entry in array *A*_1_ is a list of all the structures in Frame 2 that overlap with a particular structure in Frame 1. Similarly, *A*_2_ indicates the structures in Frame 1 that overlap with a particular structure in Frame 2.

The information in *A*_1_ and *A*_2_ allow us to determine which structures in Frame 1 are fusion candidates, and which structures in Frame 2 are fission candidates using an algorithm we call *back-and-forth structure matching*. For each structure in Frame 1, *A*_1_ is used to determine all associated structures in Frame 2. For each of these associated structures in Frame 2, *A*_2_ is used to determine all the associated structures in Frame 1. This list of Frame 1 structures comprises the fusion candidates for the Frame 1 structure being considered. An identical but opposite procedure is used to determine the fission candidates in Frame 2. The resulting two arrays of lists are denoted *W*_1_ and *W*_2_. In the remainder of automatic image analysis step we will determine which of these candidates truly underwent the mitochondrial event. The back-and-forth structure matching algorithm is illustrated in Fig 3 and is practically demonstrated with the help of a synthetic example in the supporting information.

**Fig 3.**
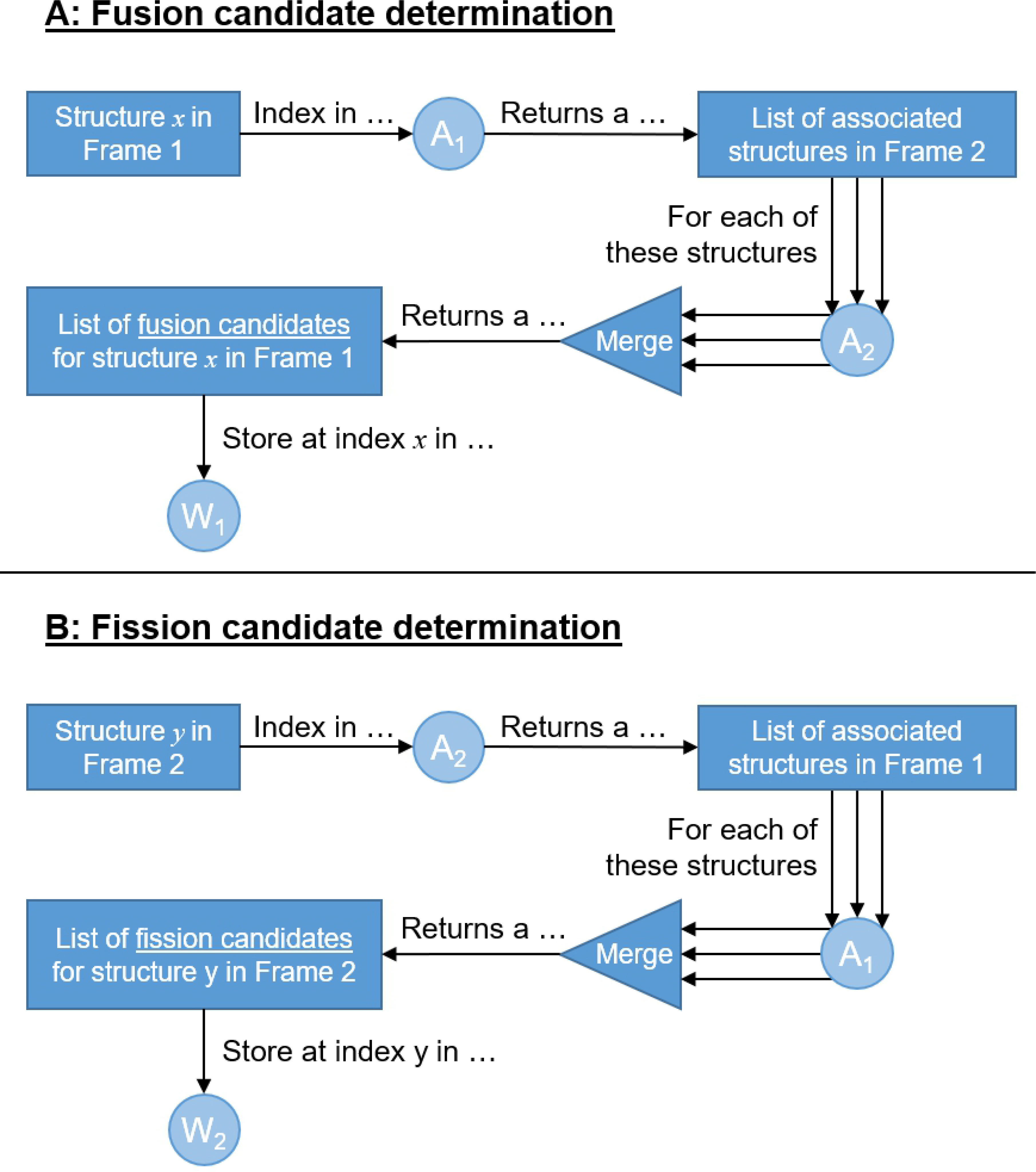
**The back-and-forth structure matching algorithm** used to determine mitochondrial structures that are candidates for fission and fusion.

Many of the mitochondrial event candidates identified in the previous step might be a result of coincidental structure overlap. These overlaps could either be due to mitochondrial movement in the time interval between Frame 1 and Frame 2, or as a result of comparing the blurred frames 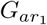 and 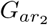. Blurring was introduced to compensate for mitochondrial movement, but has the consequence of increasing the number of false overlaps. To remove these coincidental overlaps, we begin by calculating the relative percentage overlap between associated structures in Frame 1 and Frame 2. For each structure in Frame 1, the set of associated structures in Frame 2 is identified using *A*_1_. Then, using *V*, the overlapping volume of each of these associated structures is normalised relative to the combined overlapping volume of all the associated structures. This results in the relative percentage overlap that each structure in Frame 2 has with the structure in Frame 1. In this way structures with a small percentage overlap can be considered coincidental and ignored as candidates when determining the type of mitochondrial event. These overlap percentages are stored in an array of lists *P*_1_, which has the same structure as *A*_1_. An analogous procedure is followed for all the structures in Frame 2, in this case using *A*_2_, resulting in *P*_2_. *P*_1_ and *P*_2_ are used to remove mitochondrial event candidates whose proportion of volume overlap is small.

The back-and-forth structure matching algorithm can also produce false matches. For example, when two small structures are on opposite sides of a larger central structure in Frame 1 and both fuse with this central structure in Frame 2, the smaller structures in Frame 1 will be identified as being associated with each other. This is, however, a false match since no mitochondrial event occurs between them. Such false matches can easily be avoided by detecting the presence of a third mitochondrial structure between the two candidates for association, or by determining that the candidate structures are too far apart for a mitochondrial event to be feasible. We therefore calculate the shortest vector length as well as the point in 3D space midway between all structure pairs identified in *W*_1_ and *W*_2_. This midway point represents a likely location of fission and fusion events. The shortest vector lengths can be computed from the information in 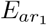 and 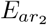. The resulting list of shortest distances are denoted by *D*_1_ and *D*_2_ and the midway points by *M*_1_ and *M*_2_ for fusion and fission respectively. Note that the length of each array in the lists matches those of *W*_1_ and *W*_2_ for the same subscript.

Using *A*_1_ and *A*_2_, we can find the structures in Frame 1 that have no associated structures in Frame 2, indicating that they will depolarise. Next, the false fission and fusion candidate structures in *W*_1_ and *W*_2_ can be filtered out by using *D*_1_, *D*_2_, *M*_1_ and *M*_2_ to determine the final sets of fission and fusion events. First, a distance threshold corresponding to the maximum distance that a mitochondrial structure could be expected to move is determined. Then, all candidates in *W*_1_ and *W*_2_ for which the respective distances in *D*_1_ and *D*_2_ exceed this threshold are removed. Next, fission and fusion candidates separated by a third structure are removed by considering the binarised stacks *B*_1_ and *B*_2_ in conjunction with the midway points in *M*_1_ and *M*_2_. Finally, by using *P*_1_, we consider two structures within Frame 1 that are both associated with the same structure in Frame 2 with a high relative percentage overlap as a fusion event. Similarly, using *P*_2_, we consider two structures within Frame 2 that are both associated with the same structure in Frame 1 with a high relative percentage overlap as a fission event. Any structure pairs with a low and high, or a low and low, combination of relative percentage overlap is ignored. Empirically we have observed that in most cases the relative percentage overlap is either above 90% or below 10%. Therefore, we consider any percentage above 50% to be high. This step results in two more arrays of lists. The *Fusion* array indicates which structures in Frame 1 will fuse, depolarise, or undergo no event. The *Fission* array indicates which structures in Frame 2 are the result of a fission event.

The final step is to generate an output image by superimposing a colour label, in three-dimensional space, on the z-stack to indicate the location of each mitochondrial event at the midway point between the two structures that fuse in Frame 1, or using the midway point between the two structures that are the result of fission in Frame 2 as the location at which a structure in Frame 1 will undergo fission. The structures that will depolarise are indicated by placing a colour label at its centre of mass, which is calculated from the z-stack for the structure in question from the 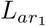 array.

The automatic image analysis step, indicating which stacks and list of arrays are used for which parts of the algorithm, is shown in Fig 2. The tuneable parameters in this step is the volume overlap threshold, the percentage that is the threshold between a low and high relative overlap percentages and the distance threshold between structures that can be considered to be related to each other. In order to further clarify each step in the MEL procedure, we have included its application to a simple synthetic sample as a supplementary section.

### Biological sample investigation

Using a mammalian cell model, we apply and validate MEL by assessing the mitochondrial network under physiological as well as disrupted conditions. For this we analysed two control cells and contrast these with cells treated with hydroxychloroquine sulphate (HCQ), leading to mitochondrial dysfunction [15].

### Cell Line Maintenance

U-118MG cells were purchased from the American Type Culture Collection (ATCC) and supplemented with Dulbecco’s Modified Eagles Medium (DMEM), 1% penicillin/streptomycin (PenStrep) (Life Technologies, 41965062 and 15140122) and 10% foetal bovine serum (FBS) (Scientific Group, BC/50615-HI) and incubated in a humidified incubator (SL SHEL LAB CO_2_ Humidified Incubator) in the presence of 5% CO_2_ at 37°C.

### Microscopy

Live cell confocal microscopy of mitochondrial fission and fusion events was conducted using a Carl Zeiss Confocal Elyra S1 microscope with LSM 780 technology. Prior to imaging, U-118MG cells were seeded in Nunc® Lab-Tek® II 8 chamber dishes and incubated with 800nM tetramethylrhodamine-ethyl ester to allow for the visualisation of the mitochondrial network (TMRE, Sigma Aldrich, 87917) for 30 minutes in the presence of 5% CO_2_ at 37°C. In order to perturb the mitochondrial network, cells were exposed to 1mM hydroxychloroquine sulphate (HCQ) (Life Technologies, T669). HCQ has previously been shown to result in fragmentation of the mitochondrial network by disrupting electron transport chain efficiency [15].

For the two control samples the time interval between frames were 6s and 20s respectively. For the HCQ treated samples the time interval was 61.7s.

### Visualising the confidence of mitochondrial events

The accuracy of the results that are produced with MEL can be affected by image quality. Therefore, to compensate for the variable image quality of the biological samples, we used MEL to calculate the position of all mitochondrial events for three different contrast stretching parameters in the pre-processing phase. This leads to three output images, each with markers indicating the location of all mitochondrial events. The output images were then divided into non-overlapping 10-by-10-by-2 voxel blocks (x, y and z-axes respectively).

Within each such block, the number of times each type of mitochondrial event was detected was counted for the three different pre-processing parameters. We then visualise the different types of mitochondrial events using different colour labels, where the intensity of these labels is proportional to the relative frequency with which each type of mitochondrial event was detected within each 10-by-10-by-2 voxel block. The intensity therefore serves as an indication of how confident MEL is that the mitochondrial event is occurring at that location.

## Results

The ability to automatically locate and count the number of mitochondrial events that occur in a sample is key to the investigation of mitochondrial dynamics. In this section, we assess and validate MEL by applying it to four different time-lapse sequences of mammalian cells that represent a range of example cases.

Figs 4 and 5 demonstrate the application of MEL to two control time-lapse sequences of cells under physiological conditions and Figs 6 and 7 consider the application of MEL to two cells that were treated with HCQ. The control cells were acquired with relatively short time intervals between the image frames, allowing individual mitochondrial structures to be more easily matched between consecutive frames, while the treated cells were acquired with larger time intervals between the image frames. This was done to validate the ability of MEL to detect mitochondrial events in conditions characterised by decreased spatio-temporal resolution.

**Fig 4.**
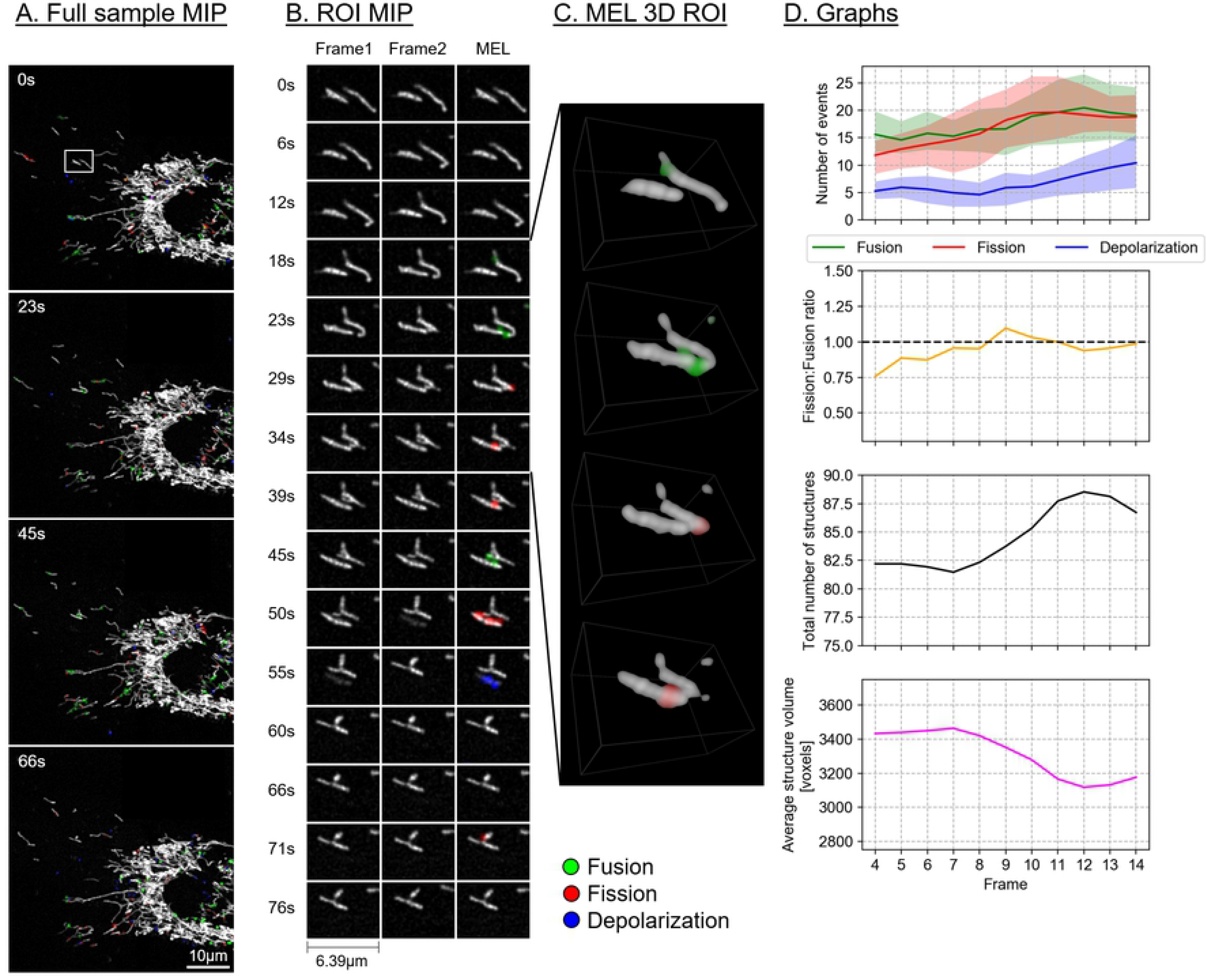
Control sample 1. A. Maximum intensity projection (MIP) of the MEL events overlaid on the entire sample image of every fourth frame in the time-lapse sequence. B. A region of interest (ROI) selection, indicated by the white square in column A. Frame 1 matches the time indicated, Frame 2 shows the subsequent time step, MEL shows the detected mitochondrial events overlaid on Frame 1. C. A selection of the ROI frames in column B, visualised in 3D using volume rendering. D. The five-frame simple moving average of the values calculated by MEL. For the number of events, the area plots indicate the maximum and the minimum that MEL detected for different pre-processing parameters.

**Fig 5.**
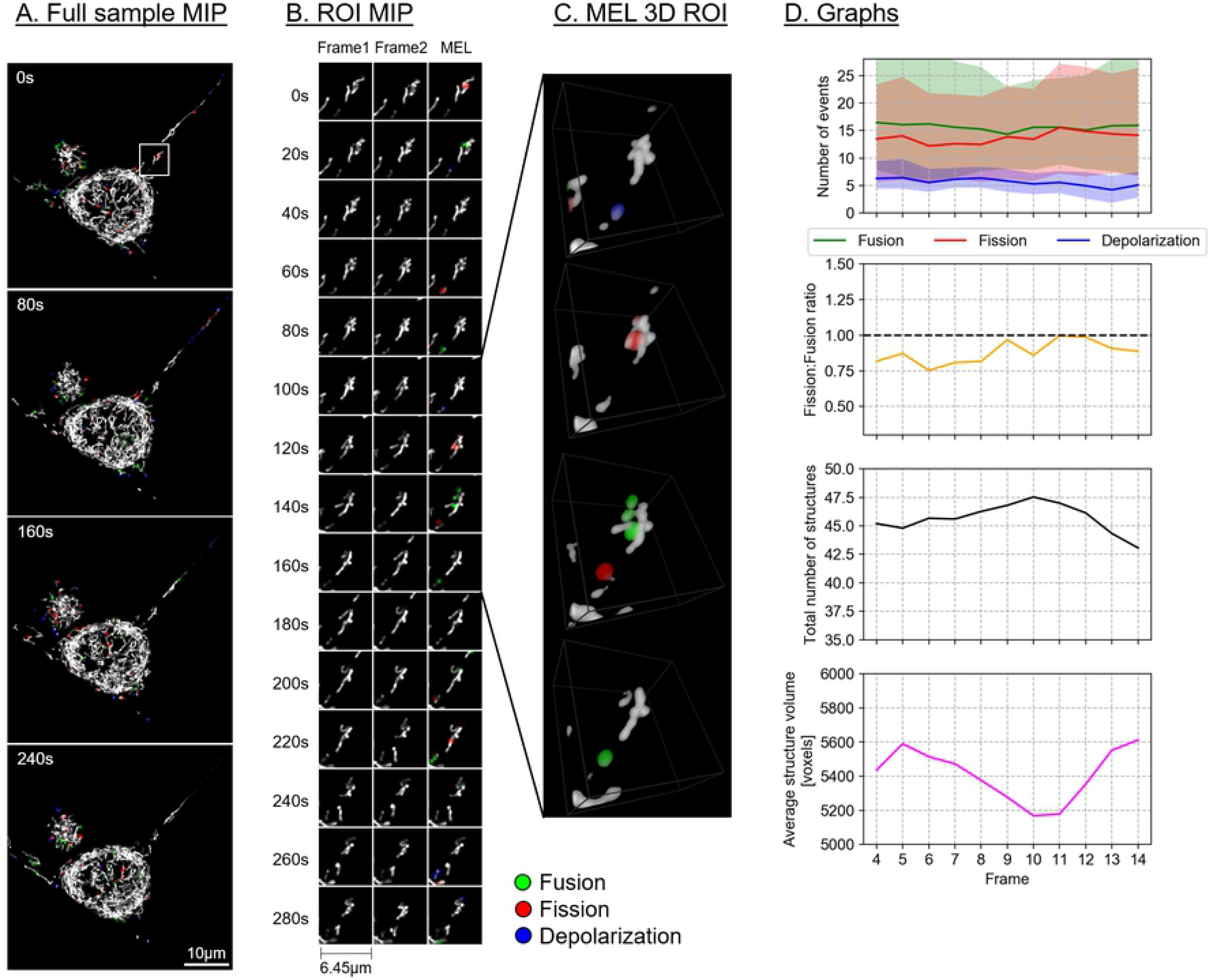
Control sample 2. Description as for Fig 4.

**Fig 6.**
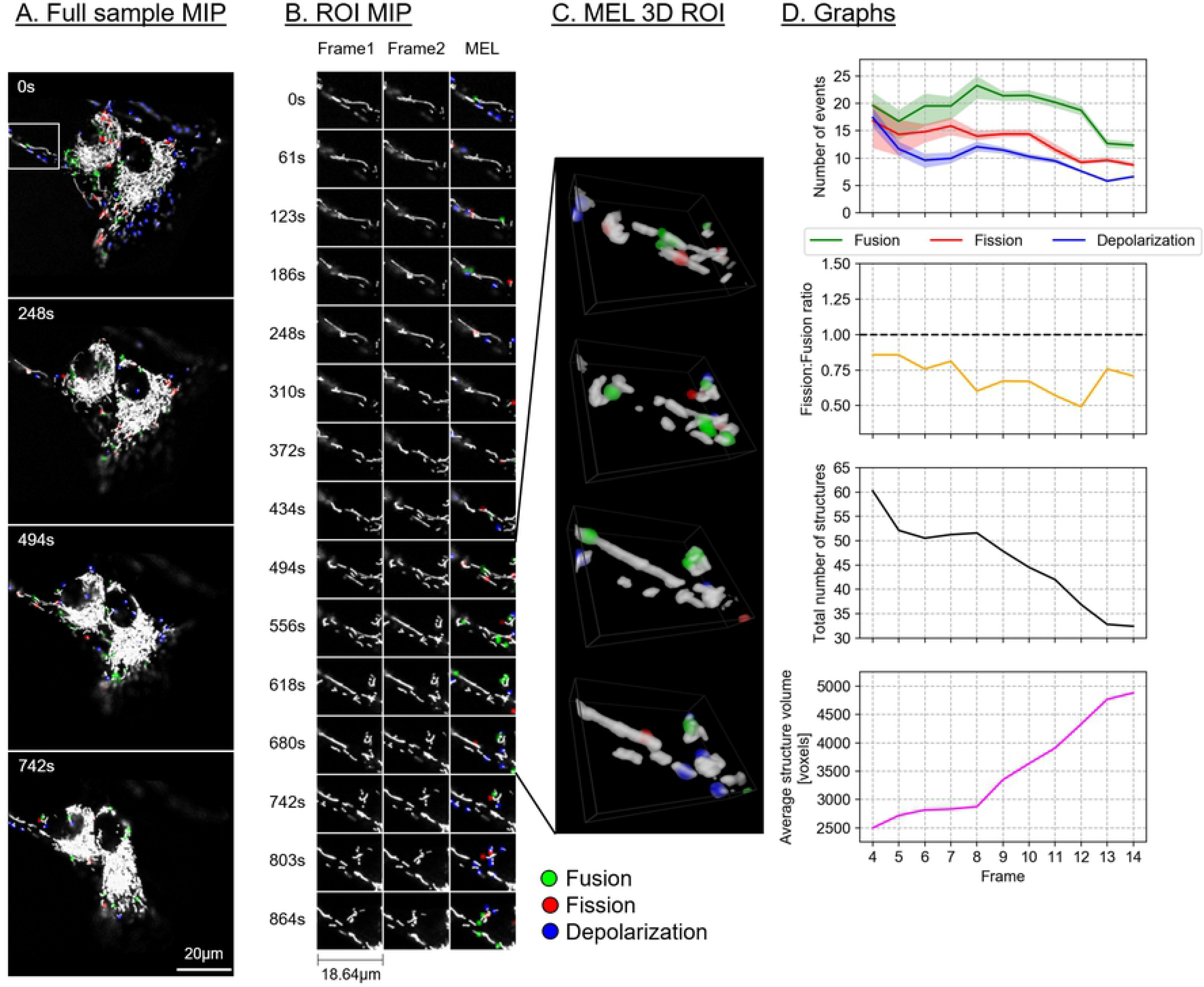
HCQ treated sample 1. Description as for Fig 4.

**Fig 7.**
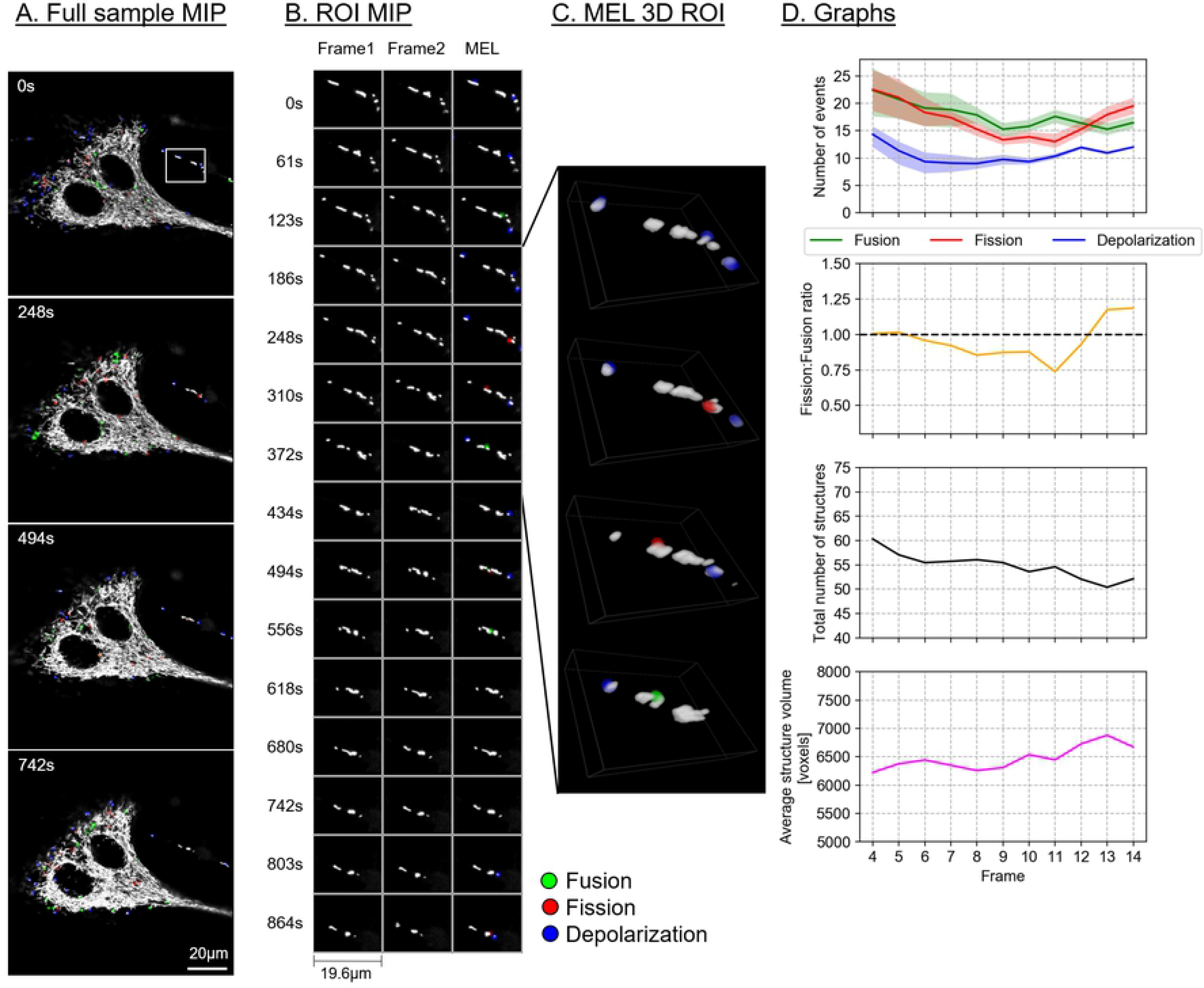
HCQ treated sample 2. Description as for Fig 4.

Figs 4–7A show the maximum intensity projection (MIP) of the MEL output for every fourth frame in the time-lapse sequence. Column B shows the MIP of a region of interest (ROI), indicated by a white square in column A, for all the frames in the time-lapse sequence. The ROI was specifically chosen to highlight the detection and visualisation of mitochondrial events. This column shows both Frame 1 and Frame 2 that was used by MEL as well as the generated output frame. Column C shows a selection of the MEL ROI frames in three dimensions. Column D shows a graphical summary of some of the data extracted by MEL for each frame pair. This includes the five-frame simple moving average (SMA) of the number of mitochondrial events that occur, the fission/fusion ratio, the total number of mitochondrial structures, as well as the average volume of these structures. Lastly, Fig 8 shows the 3D visualisation of the four cells with the mitochondrial event locations overlaid on the mitochondrial network. All sample visualisation and ROI selection was accomplished using a virtual reality based biological sample processing system [23, 24]. The distribution of the mitochondrial events throughout the sample can only be fully appreciated when visualised in virtual reality since this provides an unambiguous interpretation of the location of the events within the complex mitochondrial network.

**Fig 8.**
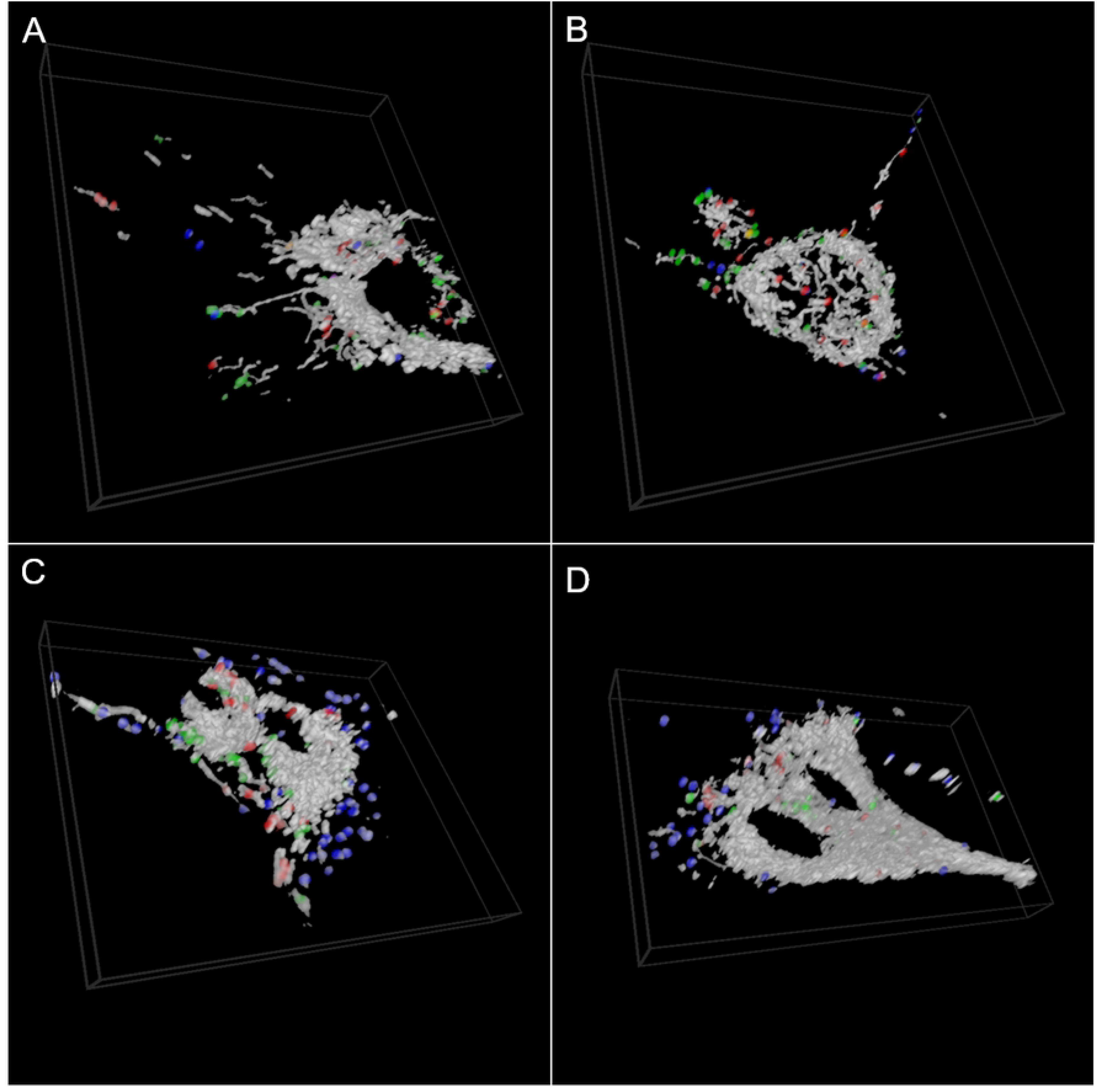
**3D visualisation of MEL event** showing the first frame in the time-lapse sequence in for the Figs 4-7. A. Control sample 1. B. Control sample 2. C. HCQ treated sample 1. D. HCQ treated sample 2.

Finally, the average percentage of structures that will undergo fission, fusion and depolarisation in every fourth frame in the time-lapse sequence (shown in column A of Figs 4-7) was calculated, and the results are summarised in Table 1.

**Table 1.**
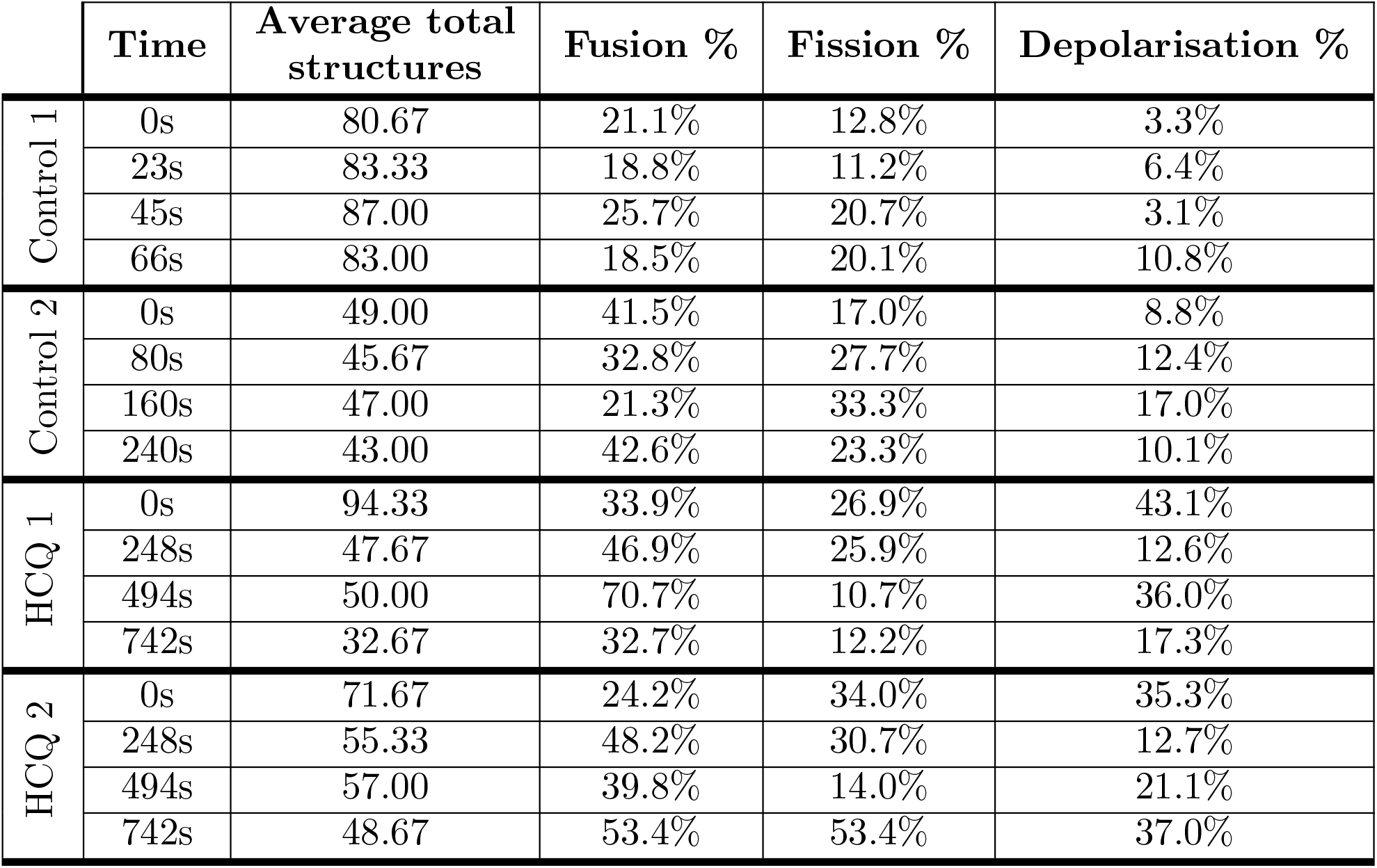
**Mitochondrial event percentages** for two control samples and two cells treated with HCQ at four different time points corresponding to column A in Figs 4-7.

|  | Time | Average total structures | Fusion % | Fission % | Depolarisation % |
| --- | --- | --- | --- | --- | --- |
| Control 1 | 0s | 80.67 | 21.1% | 12.8% | 3.3% |
|  | 23s | 83.33 | 18.8% | 11.2% | 6.4% |
|  | 45s | 87.00 | 25.7% | 20.7% | 3.1% |
|  | 66s | 83.00 | 18.5% | 20.1% | 10.8% |
| Control 2 | 0s | 49.00 | 41.5% | 17.0% | 8.8% |
|  | 80s | 45.67 | 32.8% | 27.7% | 12.4% |
|  | 160s | 47.00 | 21.3% | 33.3% | 17.0% |
|  | 240s | 43.00 | 42.6% | 23.3% | 10.1% |
| HCQ 1 | 0s | 94.33 | 33.9% | 26.9% | 43.1% |
|  | 248s | 47.67 | 46.9% | 25.9% | 12.6% |
|  | 494s | 50.00 | 70.7% | 10.7% | 36.0% |
|  | 742s | 32.67 | 32.7% | 12.2% | 17.3% |
| HCQ 2 | 0s | 71.67 | 24.2% | 34.0% | 35.3% |
|  | 248s | 55.33 | 48.2% | 30.7% | 12.7% |
|  | 494s | 57.00 | 39.8% | 14.0% | 21.1% |
|  | 742s | 48.67 | 53.4% | 53.4% | 37.0% |

## Discussion

We have described a novel method, MEL, which localises the key mitochondrial fission, fusion and depolarisation events based on a time-lapse image sequence. We provide evidence that, when using MEL, the relationship between fission and fusion can be described quantitatively. This allows the assessment of whether a cellular system is in mitochondrial fission/fusion equilibrium and whether mitochondrial depolarisation events are kept within a physiological range. We also visualise where and when mitochondrial events, specifically fission, fusion and depolarisation, take place in the cell. Given the regional complexity of cells, which may distinctively change in disease, with a particular distribution profile of organelles, mitochondria, ATP generation and vesicle dynamics associated with mitochondrial quality control, MEL may be used to dissect localised regions of dysfunction or molecular defect precisely [25–29]. Moreover, when applied in scenarios where particularly mitochondrial pathology is implicated, such as in neurodegeneration or mitophagy, precise mitochondrial event localisation as provided by MEL may be of benefit [30]. Finally, due to the relatively simple data generation approach, a role for diagnostics, especially in the context of high-throughput imaging approaches, may be envisaged [31].

MEL allows the automatic identification and subsequent visualisation of mitochondrial events over the time course of sample acquisition. This allows the dynamics of such mitochondrial activity to be studied. Data extracted by applying MEL, as shown in column D in Figs 4-7 and Table 1, reveal that the control cells are characterised by the maintenance of a fission/fusion equilibrium, with limited occurrence of depolarisation events throughout the time-lapse sequence. In contrast, HCQ treated cells exhibit much greater fluctuations in the fission/fusion equilibrium as well as a greater extent of depolarisation.

From a morphological point of view, we observe a widely spread, highly interconnected mitochondrial network for the control samples (column A in Figs 4 and 5). We note that the peripheral cell regions are characterised by a loosely defined network morphology, with few denser structures that are localised more centrally. We further notice that both fission and fusion events localise in both central and peripheral regions, in what appears to be a random fashion, with no particular focus regions or hotspots of occurrence. In contrast, the HCQ-treated samples display a very different and distinct event profile for all mitochondrial events. Most notably, a strong occurrence of depolarisation (blue) was localised primarily in the peripheral regions, a pattern that is maintained throughout the time of acquisition (Figs 6 and 7). A bias toward fusion events (green) was also seen as the network morphology becomes more densely networked, indicating mitochondrial injury. This results in fewer mitochondrial structures, each with a higher average volume. This is, to our knowledge, the first time that depolarisation events could be localised in this manner and precision, within the context of fission and fusion events.

In general, there is very little change in the overall mitochondrial network pattern of the control samples (column A in Figs 4 and 5), even when taking the shorter interval between samples into account. In comparison, the mitochondrial network of the HCQ-treated samples (column A in Figs 6 and 7) initially appears more loosely networked and, as time progresses, is seen to become denser and centred with swollen mitochondrial morphology. This accounts for the decrease in the mitochondrial events, as well as the more consistent results for different pre-processing parameters in later frames of the time-lapse sequence.

### Fission and fusion events

Most the fission and fusion events were observed around the edges of the mitochondrial network, with very few events detected in the strongly networked areas. Previous work has suggested that the density of the mitochondrial network influences fission and fusion events [16].

Using Fig 4B and C as an example, it was often observed that the same structures alternate between fission and fusion events (between 23-50s), with possible eventual depolarisation of one of the structures (55s). This scenario is supported by previous work [32, 33], and is sometimes referred to as ‘kiss-and-run’, albeit without the 3D quantitative approach and the depolarisation context.

### Depolarisation events

To our knowledge, MEL is the first approach that allows the automatic localisation of depolarisation events. This is specifically significant since depolarisation of the entire mitochondrial network has been associated with caspase-3 activation, the execution of apoptotic cell death [34] and generally demarcates the point of no return (PONR) for apoptosis [35]. Therefore, MEL may be of particular value in the quantitative assessment of cell death onset, associated with a wide range of diseases.

Depolarisation was observed to be most abundant in the HCQ treated samples, with the majority of the depolarisation events localised in mitochondria that become isolated from the main mitochondrial network and have a small average volume. This was usually observed to occur in the cell periphery (column A in Figs 6 and 7).

### Limitations of MEL

The samples that were investigated here represent a variety of intervals between image frames as well as resolutions. From the analysis it seems that MEL is robust to these variations when the movement of mitochondrial structures between two frames is small enough.

One limitation of using binarised images is that two mitochondria passing each other, due to the motility of mitochondria, will be labelled as a fusion event at one point in the time-lapse and as a fission event at a later point [16]. This is a limitation of the resolving power of the microscope used, however. Since our analysis is based on 3D z-stack data, the false detection of these events is reduced.

MEL does have tuneable parameters that affect how it performs. These are the pre-processing parameters as well the percentage and distance thresholds, which are used to remove false matches. The tuneable parameters are summarised in Table 2. For the investigation described in our Methods, these parameters were selected based on empirical observation. While we believe that the chosen values for these parameters generalise well, an automated statistical method could also be employed to determine them based on the z-stack under analysis. Such a strategy was used, for example, for an automated image analysis algorithm that determines the optimal filtering parameters to produce the best 3D quantification of mitochondrial morphology [36].

**Table 2.** The tuneable parameters used by MEL, and the values used for experimentation.

| Tuneable parameter | Value |
| --- | --- |
| Noise volume | 20 voxels |
| Gaussian blur (x-y) | $\sigma = 1.5$ |
| Gaussian blur (z) | $\sigma = 0.5$ |
| Volume overlap threshold | $\sigma = 1\%$ |
| Relative percentage threshold | 50% |
| Distance threshold | 20px |

One challenge with MEL is that it is dependent on the quality of the binarised frames (*B*_1_ and *B*_2_). The ability of Otsu thresholding to perform well is in turn dependent on the quality of the input images. It is for this reason that we first normalise the images before applying Otsu thresholding to ensure it produces the most consistent results. However, if the input image quality is poor, a small adjustment in pre-processing parameters can have a large impact on the resulting binarised image. The consequence of this is that some structures could be joined together for some pre-processing parameter choices and be binarised as separate structures with others. The result of this can be seen for our control samples, where there is a large difference between the minimum and the maximum number of mitochondrial events for different pre-processing parameters (shown in Figs 4D and 5D). This can be ascribed to the very limited dynamic range in the input image intensity, which is a consequence of the acquisition parameters that were chosen to ensure that longer time-lapses in 3D can be acquired while limiting the amount of bleaching and phototoxicity. Hence, well controlled acquisition parameters and an awareness of the dynamic range will benefit the application of MEL.

## Conclusion

Mitochondrial events, particularly fission, fusion and depolarisation, play a central role in cellular homeostasis, function and viability. However, the precise localisation of these events within a cell as well as the quantitative characterisation thereof remains challenging.

We have presented MEL (mitochondrial event localiser), an approach we have developed to automatically determine the three-dimensional location of fission, fusion and depolarisation events in a sample from a three-dimensional time-lapse sequence. The number of mitochondrial structures, their combined and average volume, as well as the number of events at each frame in the time-lapse sequence can subsequently be calculated to provide a quantitative assessment.

To demonstrate its performance, we have applied MEL to three-dimensional time-lapse sequences of two control samples and two hydroxychloroquine sulphate treated samples with varying time intervals and image resolutions. The analysis confirmed that the fission/fusion equilibrium is maintained throughout the time-lapse of the control cells, as is commonly expected for healthy cells. This is not the case for the treated samples, both of which showed an initial bias toward fusion, causing the mitochondrial network to become denser and more perinuclear localised.

It is hoped that the automatic localisation and quantification of mitochondrial events offered by MEL can support research into mitochondrial function in health and disease, which may consequently permit new diagnosis and treatment strategies to be implemented for diseases associated with mitochondrial dysfunction.

## Supporting information

### Synthetic example

In this section we demonstrate the steps of the MEL algorithm by applying it to two synthetically generated image frames. This is intended only to clarify the MEL algorithm, and does not represent a realistic scenario as would be expected when analysing mitochondrial events. Although MEL is intended for application to three-dimensional samples, we will consider two-dimensional images here for clarity.

Since our images are synthetic, we omit the normalisation step shown in Fig 1 and begin by showing binarised Frames 1 and 2 in S1 Fig. We also overlay these binarised images to make it easier to identify those structures that will fuse or undergo fission from Frame 1 to Frame 2. Next, each structure in the binarised frames is given a unique label number, with 0 indicating the background. Each labelled structure is then separated to create an array of images (not shown in S1 Fig) which is Gaussian filtered to allow for a less strict structure overlap matching between Frame 1 and Frame 2. The array of labelled images are also Canny filtered, leaving only the pixels on the edge which is used to determine the distance between structures and the location of the fission and fusion events. S1 Fig shows only the first labelled structure in the array for each frame. These images are then analysed according to the process in Fig 2 to produce a set of locations and types of mitochondrial events that are hypothesised to have occurred in the time between Frame 1 and Frame 2. The result is then overlaid with Frame 1 using colour labels for the different events.

**S1 Fig. Synthetic image example process flow.**

The matrices and arrays calculated for the synthetic example by the process depicted in Fig 2 are shown in S2 Fig. The overlap matrix *V* is calculated by multiplying all combinations of the Gaussian filtered structures, one from Frame 1 and one from Frame 2, to determine a representation of the volume. Label number 0 indicates the background and therefore is left blank throughout.

In the synthetic example shown, there is a small overlap between Frame 1 label 4 and Frame 2 label 2 due to the blur which the filter has introduced. To compensate for such coincidental matches, all overlapping volumes that account for less than 1% of the volume of either structure in question are eliminated, and is consequently shown as 0 in the table (S2 Fig).

**S2 Fig. Synthetic image example calculated matrix and list of arrays.**

From matrix *V*, the arrays *A*_1_ and *A*_2_ are determined by simply reducing *V* to indicate which structures in the other frame presented with a non-zero overlapping volume. The relative percentage overlap, *P*_1_ and *P*_2_, of each structure in one frame with all associated structures in the other frame can then be calculated from *A*_1_ and *A*_2_. This is in effect a normalisation of the volumes in matrix *V*, where the volume of a certain structure combination is divided by the total volume of the given structure in either Frame 1 (producing *P*_1_) or Frame 2 (producing *P*_2_). Each row, therefore, sums to 100%.

Using the back-and-forth structure matching described in Fig 3, we generate *W*_1_ and *W*_2_, to indicate which structures are associated with each other in the same frame. These are candidates for fission and fusion events, although some of them could be false matches that the next step aims to eliminate. It is worth noting that each structure combination occurs twice in the array of lists. Even though we show the calculations that follow for both, it is only necessary to use one.

Using *W*_1_ and *W*_2_ along with the edge array of images *E*_*ar*_ for Frame 1 and Frame 2, we find the shortest distances, *D*_1_ and *D*_2_, as well as the midway points, *M*_1_ and *M*_2_, between each combination of structures. If the shortest distance between the candidate structures is above a set threshold (in the case of this synthetic sample this was 50 pixels), the two structures are considered unrelated and is ignored in the visualisation (refer to S1 Fig and S3 Fig). Secondly, if the midway point is sampled from the binarised images, *B*_1_ and *B*_2_, and coincides with another structure it is also considered a false match. This is because two structures cannot fuse through a third separating structure. Rather, they would individually fuse with the central structure. A similar logic applies when considering fission events in Frame 2. This is illustrated in S3 Fig for a fusion, fission and unrelated label combination.

**S3 Fig. Example of determining fission and fusion event locations.**

Now the mitochondrial event status can be determined for each candidate structure combination in *W*_1_ and *W*_2_. Structures in Frame 1 are labelled “Fuse”, “Depolarize”, or “Unrelated”, while structures in Frame 2 are labelled “Fission”, or “Unrelated”.

Finally, using the midway points, as well as the centre of mass of the structures to indicate the location of the mitochondrial event, along with the status arrays, the final output image is generated (S1 Fig).

## Acknowledgments

The authors gratefully acknowledge the support of NVIDIA Corporation with the donation of the GPU used for this research. The authors acknowledge financial support from the South African National Research Foundation (NRF), the South African Medical Research Council (SAMRC), the Cancer Association of South Africa (CANSA), and Telkom South Africa. We wish to thank the Cell Imaging Unit, Central Analytical Facility (CAF), Stellenbosch University for providing technical support.

